# ALOX15, a new potential functional target of lung adenocarcinoma

**DOI:** 10.1101/2023.03.03.531049

**Authors:** Xiaocui Liu, Yangyang Tang, Hui Liu, Shupeng Zhang, hongshu Sui, wenwen Sun, Siyu Xuan, minhua Yao, ping Song, peng Qu, yanping Su

**Author notes:** Correspondence: yanping Su. These authors have contributed equally to this work and share first authorship (Xiaocui Liu, Yangyang Tang).

## Abstract

**Purpose:** The purpose of this study is to explore whether the down-regulation of ALOX15 is related to the stage, differentiation and prognosis of lung adenocarcinoma, whether the overexpression of ALOX15 can inhibit tumor proliferation and metastasis, and whether it is related to the functional target of lung adenocarcinoma(LUAD).

**Methods:** Firstly, bioinformatics of lung cancer patients was analyzed using the TCGA database to study the differential expression of ALOX15 in LUAD and its effect on the survival of LUAD. Then, 50 clinical tissue samples of LUAD were collected to detect the expression of ALOX15 and its relationship with the differentiation degree and stage of lung cancer. Finally, the relationship between the expression of ALOX15 and the proliferation and differentiation of LUAD cell lines (NCI-H1944, A549, PC9) with different degrees of differentiation and the construction of ALOX15 overexpression stable lines was detected.

**Results:** ALOX15 bioinformatics analysis showed that ALOX15 decreased significantly in the early stage of LUAD and had no correlation with the survival of lung cancer patients. ALOX15 was downregulated in LUAD with low differentiation and metastasis in LUAD. LUAD cell lines showed that the lower the degree of differentiation, the lower the expression of ALOX15, and the overexpression of ALOX15 in LUAD cells inhibited the proliferation and migration of cancer cells.

**Conclusion:** These results suggest that the expression of ALOX15 is closely related to the differentiation, proliferation, and metastasis of LUAD, and that upregulation of ALOX15 may inhibit the development of LUAD, suggesting that ALOX15 is a potential biological therapeutic target.

**Author summary:** *Why was this study done?:*

- Lung cancer is one of the most frequently diagnosed cancers in the world and the leading cause of cancer-related deaths worldwide.
- Lung cancer is a heterogeneous disease with a wide range of clinicopathologic features. Lung cancer is roughly divided into non-small cell lung cancer (85% of all diagnoses) and small cell lung cancer (15% of all diagnoses). Adenocarcinoma is a common subtype of non-small cell lung cancer, and its recurrence rate is high, and the prognosis is poor. Therefore, the pathogenesis and characteristics of adenocarcinoma are studied and explored.

*What did the researchers do and find?:*

- Biological information database was used to analyze the expression level of arachidonic acid-15-lipoxygenase (ALOX15) in lung adenocarcinoma, and then the expression differences were discussed through clinical samples and cell experiments.
- Low ALOX15 expression was detected in lung adenocarcinoma (LAUD) patients compared with normal tissues, and ALOX15 levels influenced LUAD development.
- By infecting A549 with lentivirus and overexpressing ALOX15 of A549 and PC-9 with PC9, it was found that ALOX15 inhibited the proliferation of tumor cells

*What do these findings mean?:*

- arachidonic acid-15-lipoxygenase may be one novel potential biomarker for LUAD and a potential therapeutic target

## Introduction

Previous results have shown that chronic irritation and chronic inflammation in the lungs of some populations develop into chronic obstructive pulmonary disease (COPD) and even lung cancer[1, 2]. Lung cancer mainly consists of two subtypes: non-small cell lung cancer (NSCLC) and small cell lung cancer (SCLC). NSCLC accounts for nearly 80 percent of lung cancer cases and includes the two main types, lung adenocarcinoma (LUAD) and lung squamous cell carcinoma. LUAD is the dominant histology, and the incidence is still increasing[3]. Despite recent advances in medical technology, most patients with lung cancer are still detected in the middle and late stages, and the 5-year overall survival rate is still less than 15%[4].LUAD can invade blood vessels and lymphatic vessels at an early stage. It often metastasizes before the primary cancer causes symptoms. The traditional treatment is ineffective. Therefore, it is particularly important to find potential biological markers for targeted therapy.

It has been reported that the development of lung cancer is closely related to chronic inflammation. Chronic pulmonary inflammation is mainly caused by change in the immune microenvironment. Mainly including the infiltration of inflammatory cells and the accumulation of cytokines, prostaglandins, and chemokines, which can stimulate a variety of processes, including cancer cell proliferation, angiogenesis, and metastasis[5]. At the same time, chronic inflammatory stimulation also causes changes in the immune microenvironment. The infiltration of inflammatory cells in chronic pulmonary inflammation are mainly caused by Myeloid derived suppressor cells (MDSCs), Tregs, M2 macrophages, and resistant dendritic cells invading the lungs[6, 7]. It can be said that the immune suppressive cells that exist in the second camp are our betrayers.

The development and maintenance of chronic inflammation is strongly influenced by oxidative metabolism of polyunsaturated fatty acids (PUFAs). The oxidative metabolism of polyunsaturated fatty acids is regulated by several groups of enzymes, among which lipoxygenase (LOXs) plays an important role[8]. Recent studies have found that arachidonic acid-15-lipoxygenase (ALOX15), as a rate-limiting enzyme of oxidable polyunsaturated fatty acids, plays an important role in the formation of key lipid mediators such as lipoproteins and dissolved proteins to end inflammation. ALOX15 has also been shown to be involved in the development of physiological and various pathological diseases, including cell maturation and differentiation, inflammatory diseases and tumorigenesis [9]. Under physiological conditions, ALOX15 is expressed in reticulocytes, eosinophils, dendritic cells, alveolar macrophages, airway epithelial cells, immature dendritic cells, vascular cells, and resident peritoneal macrophages. Its substrates are arachidonic acid (AA) and linoleic acid (LA). ALOX15 metabolizes AA to 15-hydroxyeicosapetetraenoic acid (15(S)-HETE) and 12-hydroxyeicosapetetraenoic acid (12(S)-HETE) and LA to 13-hydroxyoctadecanoenoic acid (13(S)-HODE)[1]. Metabolites are effective mediators to inhibit chronic inflammation and play an important role in inflammation-related diseases. Several reports have demonstrated the anti-inflammatory properties of ALOX15 and its metabolites [10, 11].

ALOX15 expression is also altered in a variety of cancer such as colon cancer, lung cancer, esophagus, breast cancer, endometrium, bladder and pancreatic cancer to affect cancer development [12]. ALOX15 has been implicated in many aspects of carcinogenesis, such as angiogenesis, inflammation, and metastasis, as demonstrated in solid tumors and hematological malignancies[13]. The expression level of ALOX15 is also closely related to the risk of colon cancer [10, 14]. ALOX15 acts on colon cancer through inflammatory cytokines or other pathways [15-17]. ALOX15 is lost in the early stage of colorectal cancer. re-expressed ALOX15 by plasmid and lentiviral vector can inhibit the growth of the cancer cells [10, 18, 19]. can be used as a potential biomarker for the diagnosis of colorectal cancer [15, 17]. It was found that ALOX15 inhibited peroxisome proliferator-activated receptor γ and COX-2 dependent signaling to inhibit gastric tumorigenesis [20]. But the role of ALOX15 in lung cancer is still unclear.

Because lung cancer is related to chronic inflammation[6], this study demonstrates whether the reduction of ALOX15 will lead to the development of lung cancer? Whether the continuous decrease of ALOX15 is related to the staging and metastasis of lung cancer Whether the transfection of ALOX15 can inhibit the proliferation and migration of lung cancer. The demonstration of this mechanism will provide a new target for the treatment of lung adenocarcinoma.

## Materials and Methods

### Raw data collection and processing

Clinical data of 878 patients were downloaded from TCGA database, and transcripts of 535 LUAD specimens and 59 adjacent tissues with ALOX15 gene were screened out from 878 patients. Meanwhile, we obtained the differential expression of ALOX15 in different stages of LUAD, as well as the expression of ALOX15 in TNM stages. There are three formats for RNA-seq data (COUNT, FPKM, and FPKM-UQ), and we chose the second format for further analysis. Primitive human matrices were annotated using GTF files. The Tidyverse function of R we used to match the gene expression matrix to the clinical data based on its unique ID number to generate a summary matrix for subsequent analysis.

### sample selection

We collected a total of 53 lung adenocarcinoma patients’ para-cancer tissue and cancer tissue samples, all of which were collected from archive samples and sample wax blocks of the Department of Pathology, the Second Affiliated Hospital of Shandong First Medical University, Shandong, China. The collection range of samples was from January 2020 to May 2022, and the experiment of this sample was conducted from 2021 to July 2022.

### Immunofluorescence (IF)

Fifty LUAD wax block specimens were collected (Department of Pathology, The Second Affiliated Hospital of Shandong First Medical University, Shandong Province, China). The experiment was performed according to the protocol below. After the section were cut and deparaffinized, they are washed for 3 times with PBS. To remove endogenous catalase, 3%H_2_O_2_ was added and immersed for 10min. The cells were washed 3 times with PBS for 5 min each time, then EDTA buffer was added for antigen repair, and washed as above. After cells were then blocked with 5%BSA for 1 h, primary antibody to ALOX15 (1:500, Abcam, shanghai, China) was added and incubated at 4 °C overnight. The next day, it was taken out and rewarmed at room temperature for 30min and washed 3 times with PBS for 5 min each time. Anti-rabbit fluorescent secondary antibody (1:500) was added in the dark and incubated at room temperature for 2 hours, then washed as above. Subsequently, the cells were counterstained with DAPI. The slices were sealed with an anti-fluorescence quencher. Finally, the results were observed by fluorescence microscopy.

### Cell culture

Lung adenocarcinoma cell A549 and PC-9 (Punosai, Wuhan, China) were resuscitated and cultured in Ham’s F-12K and 1640 medium containing 10% fetal bovine serum and 1% penicillin-streptomycin. The cells were placed in an incubator containing 5% CO2 at 37°C. When the degree of cell fusion reached about 90%, cell passage and subsequent experiments were carried out.

### Lentivirus infection

ALOX15 overexpressed lentiviral vector and its corresponding no-load vector were designed and constructed by OBiO Technology Company (Shanghai, China). The virus titer was 4.29×108TU/ml. It was verified by sequencing that the ALOX15 sequence in the vector was correctly inserted into the expression vector. To generate stable strains of ALOX15, A549 and PC-9 cell lines were co-transfected with lentivirus overexpressing ALOX15, and then purified with paromomycin to determine stable transfection of ALOX15 and then subculture.

### Western blot

ALOX15 protein levels in lung adenocarcinoma cell lines and clinical tissues were analyzed by western blotting. Cell lysis solution was added to extract protein and the concentration of BCA protein was determined. An equal amount of protein samples was then collected, the protein was denatured by boiling in a metal bath at 100°C for 10 minutes with the appropriate loading buffer. Separation and concentration gels were prepared using the 10%PAGE gel preparation kit. When the protein was on the glue, the voltage was 80V for 30min, and when the protein was on the separation gel, the voltage was increased to 120V for about 1h. This was followed by a constant current of 400mA for 30min. After the blocks were incubated with 5% skim milk powder blocking solution for 2h on a shaker, they were washed with TBST three times for 10min each time. Primary antibodies were added and incubated overnight at 4 ° C. Secondary antibody (1:5000) was then added and incubated at room temperature for 1 hour. ECL chemiluminescence solution was added and protein bands were scanned by chemiluminescence imaging system. Images were processed using ImageJ software to analyze the relative expression level of ALOX15 protein.

### Cell proliferation assay

The effect of ALOX15 on lung adenocarcinoma cell viability was determined using CCK-8 assay (solarbio) according to the manufacturer’s instructions. The lung adenocarcinoma cells were inoculated with 5×10^3^/well in 96-well culture plates and cultured in 37°C, 5%CO_2_ incubator for 24h. After 24 hours of culture, CCK8 reagent 10μl/well was added every 12 hours away from light and cultured in 37°C, 5%CO_2_ incubator for 1-3h. The absorbance of each sample was measured every half hour at the wavelength of 450nm.

### wound healing assays

Normal cells and cells overexpressing ALOX15 were inserted into the well plate and cultured in complete medium for 24 h. When the cell confluence was 100%, the cells were scratched with a 200μL pipette nozzle, and the well plate was washed with PBS to remove any debris and complete culture medium was added. The well plate was photographed immediately afterwards and every 12 hours for the next 36-72 hours, processing the images with the ImageJ software.

### Statistical analysis

Results were expressed as Mean ± SD. Statistical analysis was performed using one-way ANOVA combined with Student t test, and the result differences were regarded as significant at P < 0.05. Experiments were repeated independently at least three times with at least three technical replicates.

## Results

### Low expression of ALOX15 protein in LUAD in the early stage of LUAD

In this study, database statistical analysis showed that the expression of ALOX15 in LUAD samples was lower than that in normal lung tissues (Fig1.A). The expression levels of ALOX15 were further monitored in the pathological stage I-IV of lung adenocarcinoma. Compared with those expression levels in stage I LUAD, their expression level in stage II was downregulated and statistically significant (Fig1.B). At the same time, it was found that ALOX15 tended to decrease with the degree of cancer progression. In tumor TNM staging, the expression of ALOX15 is downregulated in LUAD. In the primary tumor (T), the expression of ALOX15 was significantly down-regulated (F1.C), and the difference was statistically significant. In regional lymph node involvement (N), ALOX15 expression was also significantly down-regulated (Fig1.D), and in distant metastatic LUAD (M), ALOX15 expression also tended to be down-regulated (Fig1.E). However, ALOX15 expression levels were not involved in the survival of LUAD (Fig1.F). Therefore, the expression of ALOX15 is downregulated in LUAD and influences the early differentiation of LUAD.

**Fig 1.**
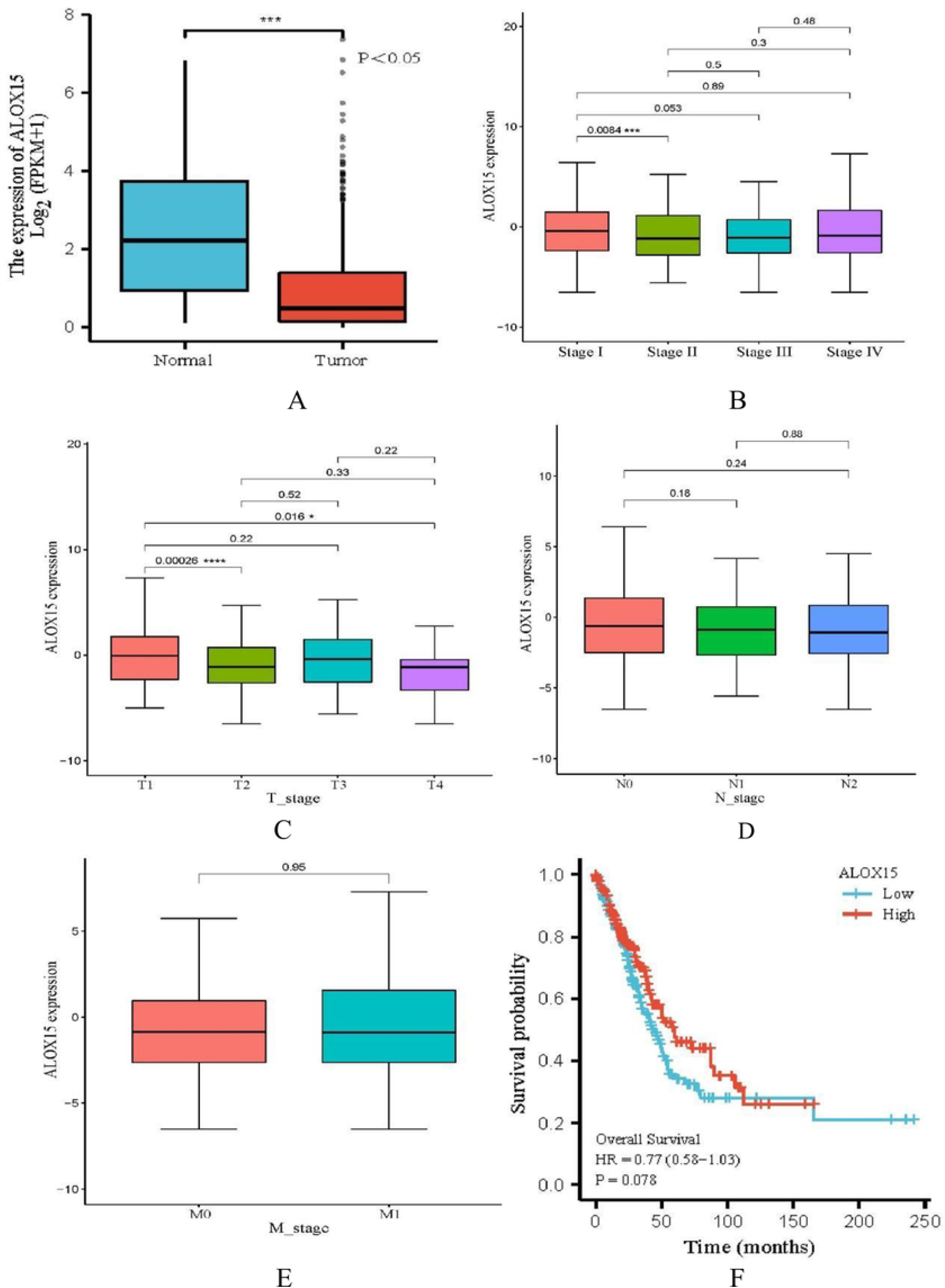
ALOX15 expression level in LUAD. (A). Differences in expression levels of ALOX15 in lung adenocarcinoma (LUAD) and normal tissues;(B). Expression of ALOX15 in clinical stage 4 LUAD. (C). Expression of ALOX15 in T stage of LUAD; (D)Expression of ALOX15 in N stage of LUAD (E). Expression of ALOX15 in M stage of LUAD. (F). Survival analysis diagram. *, P<0.05, **, P<0.01. T: primary tumor; N: Regional lymph node involvement; M: Neoplasm Distant metastasis

### The expression level of ALOX15 was related to the differentiation degree of LUAD

#### ALOX15 expression was decreased in LUAD and expressed in the nucleus

Based on the results of biology on database analysis, we collected clinical samples of LUAD to detect the expression level of ALOX15. Firstly, most of the adjacent tissues were found to have normal alveolar and bronchial structures by HE staining. But the tumor tissue lost its normal structure, the number of cells increased, and the atypia was obvious (Fig2. AB). Secondly, the expression level of ALOX15 was detected in LUAD. ALOX15 was expressed in the cell nucleus and was more expressed in the adjacent tissues of LUAD, but less expressed in the cancer tissues (Fig2.CD).

**Fig 2.**
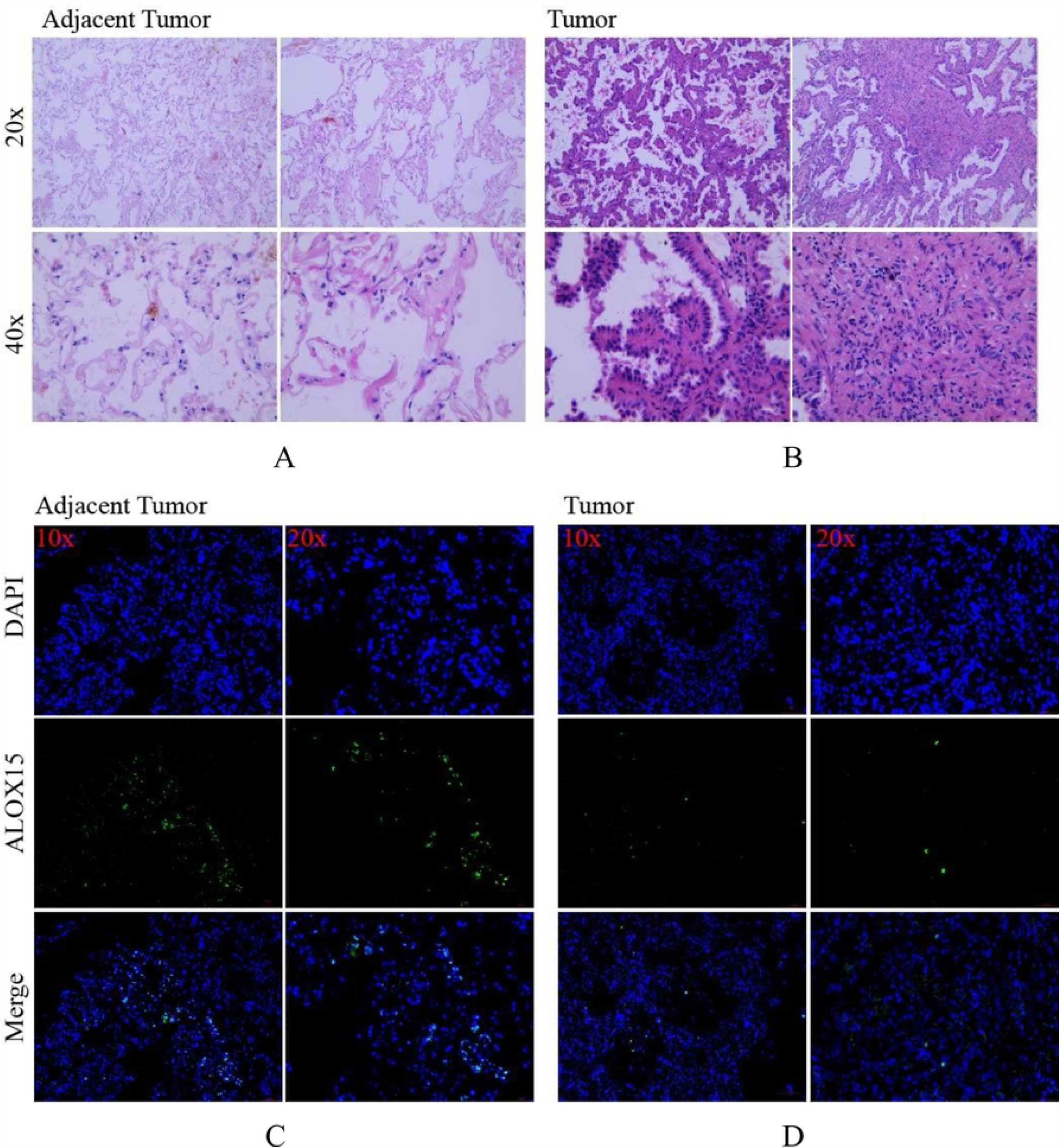
Expression of ALOX15 in lung adenocarcinoma samples. (A) (B). HE staining test found that Most of the adjacent tissues also had normal alveolar and bronchial structures. However, the normal structure of cancer tissue was lost, and the number of cells increased with obvious atypia; (C)(D) immune-fluorescence results demonstrated that the expression level of ALOX15 was different in para-cancer tissues compared with cancer tissues. The expression of ALOX15 in para-cancer tissues was higher than that in cancer tissues. *, P<0.05, **, P<0.01.

#### ALOX15 is significantly downregulated in LUAD with metastasis

For LUAD with or without lymph node metastasis, ALOX15 expression was significantly downregulated in tissues with metastasis. The expression of ALOX15 in LUAD adjacent tissues was higher than that in LUAD tissues, with significant. (Fig3. AB). Meanwhile, ALOX15 tends to decrease with the progression of LUAD.ALOX15 was downregulated more significantly in metastatic cancer than in non-metastatic cancer, with significant (Fig3.CD). Therefore, ALOX15 can be used as a biomarker for LUAD metastasis. (Table1.2)

**Fig3.**
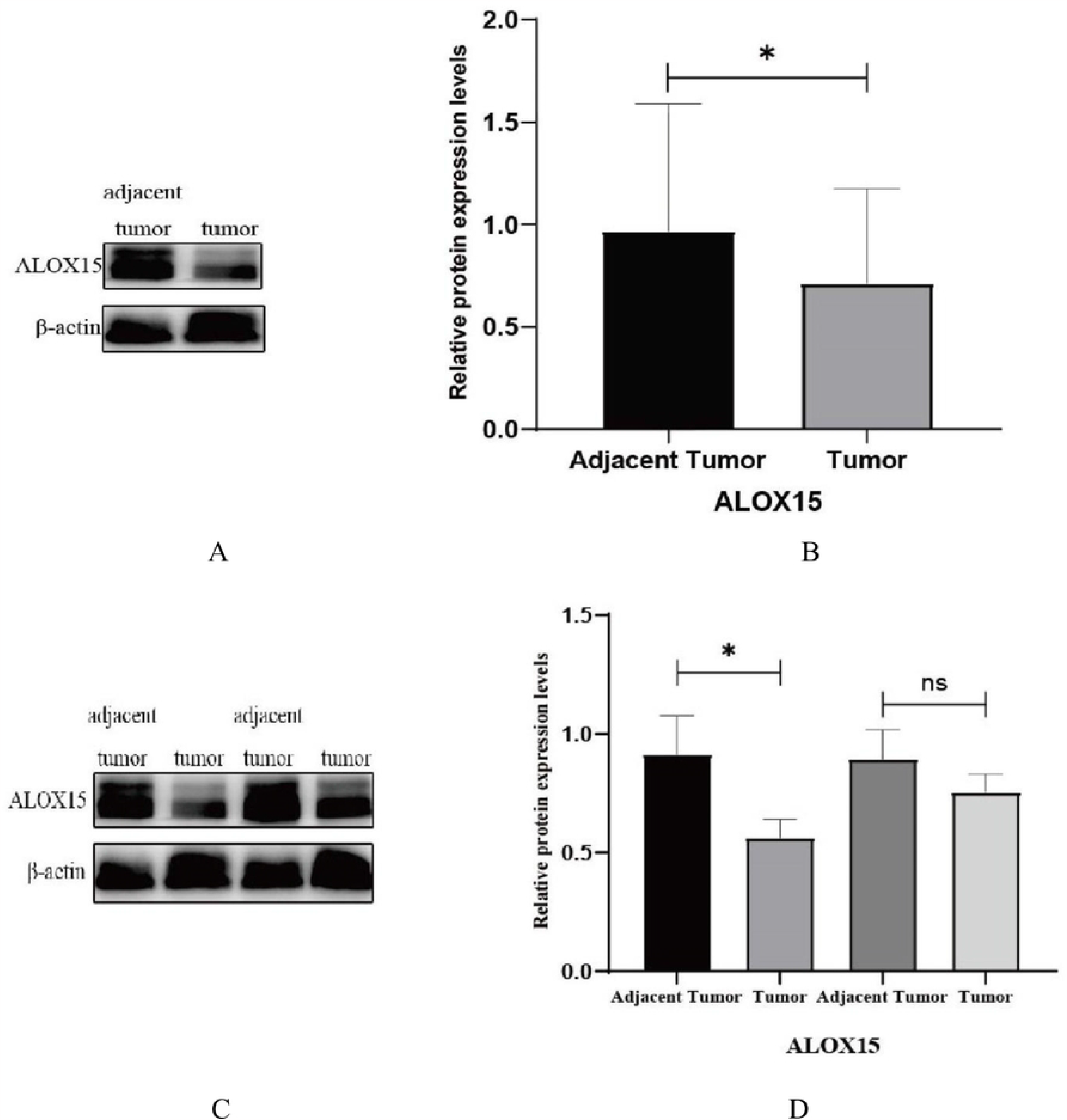
expression of ALOX15 in LUAD. (A)(B). The expression of ALOX15 in LUAD adjacent tissues and cancer tissues is shown. (C) (D). ALOX15 expression was detected in patients with and without lymph node metastasis, and ALOX15 expression was downregulated in patients with lymph node metastasis; *, P<0.05, **, P<0.01.

**Table 1.**
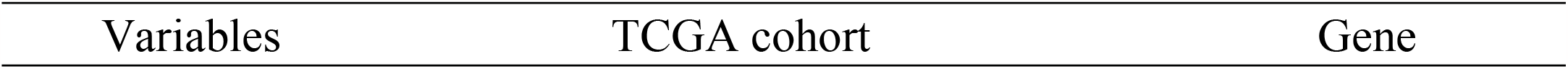
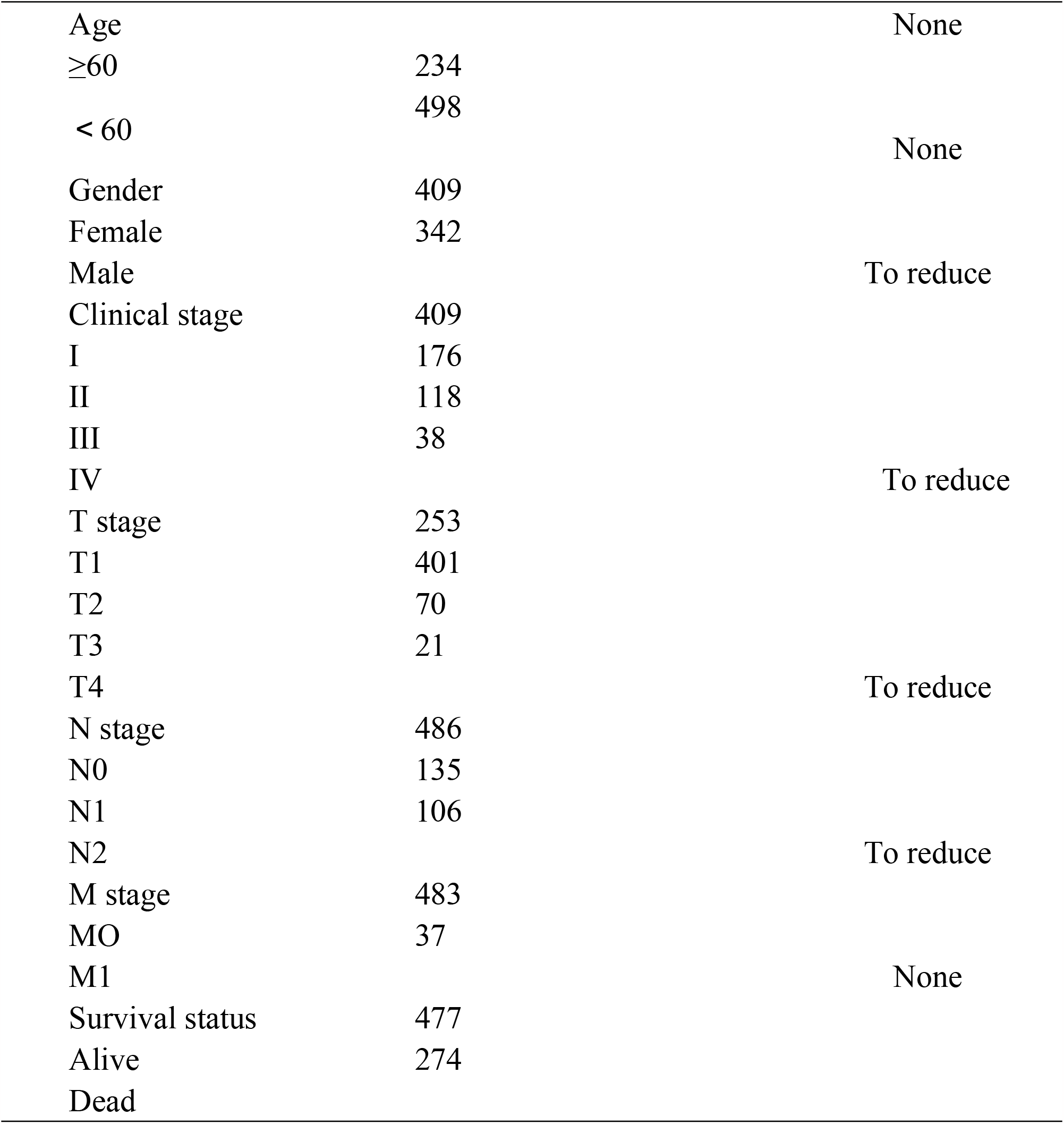
The clinical characteristics of patients and ALOX15 Gene in the TCGA.

**Table2.**
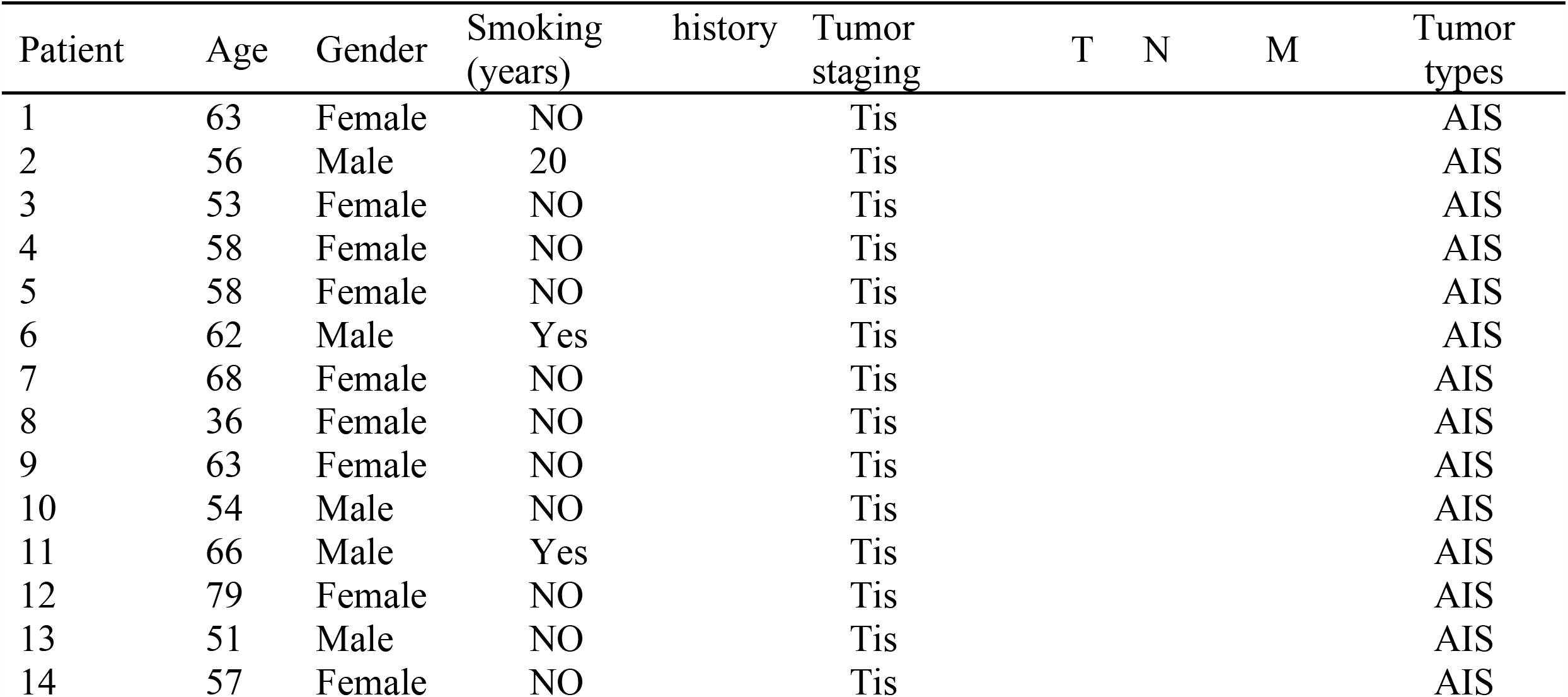
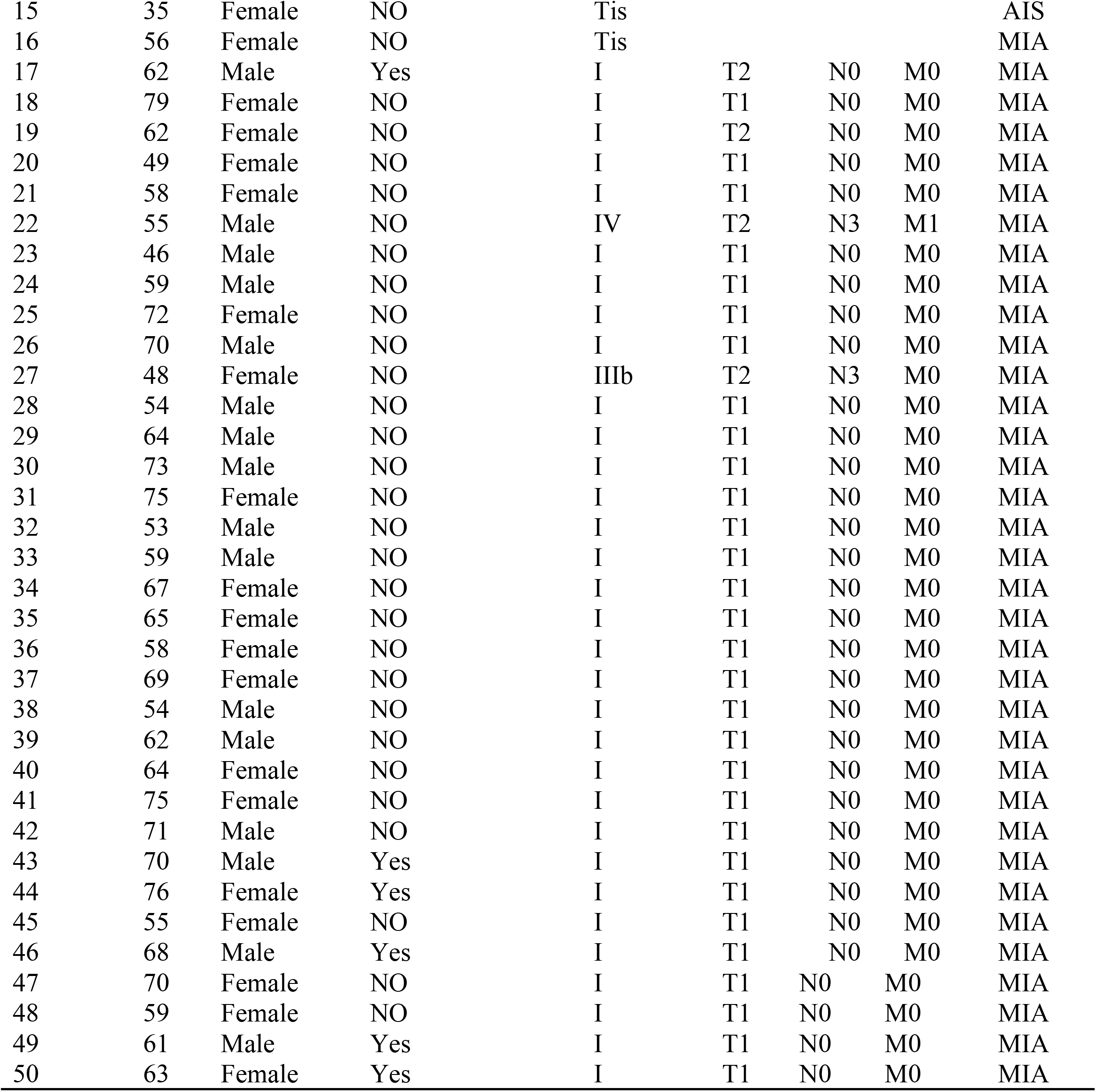
Characteristics of 50 clinical data samples.

#### The expression of ALOX15 is associated with the differentiation of lung cancer cell lines

ALOX15 was expressed in LUAD cell lines, with statistical significance (Fig4. A). The expression level of ALOX15 was found to be different in three different LUAD cell lines due to tumor differentiation. ALOX15 was highly expressed in well-differentiated adenocarcinoma PC-9, low in moderately well-differentiated adenocarcinoma A549, and much low in poorly differentiated adenocarcinoma NCI-H1944. The expression level of ALOX15 was PC-9 > A549 > NCI - H1944(Fig4. BC). Therefore, the expression level of ALOX15 in LUAD depends on the degree of differentiation of LUAD (Table 3).

**Fig 4.**
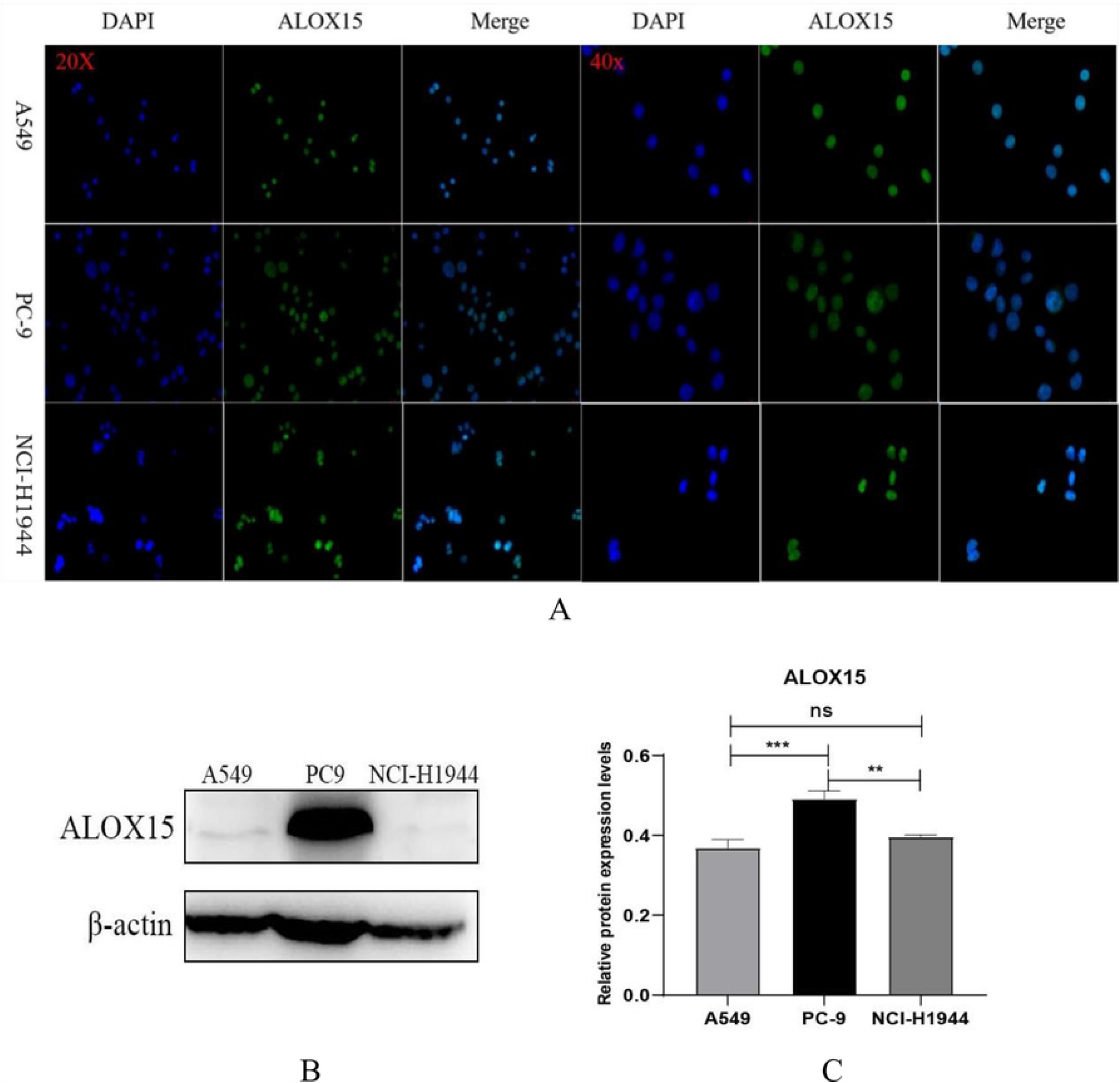
Expression of ALOX15 in different differentiated cells. (A)(B).IF results were obtained, ALOX15 is expressed in LUAD cells A549 and PC-9, At the same time, it is expressed in the nucleus, which is consistent with the expression location in tissues subsequently; (C)(D).Western blot results showed that the expression levels of ALOX15 were different in the two LUAD cell lines (A549, PC-9). *, P<0.05, **, P<0.01.

**Table3.**
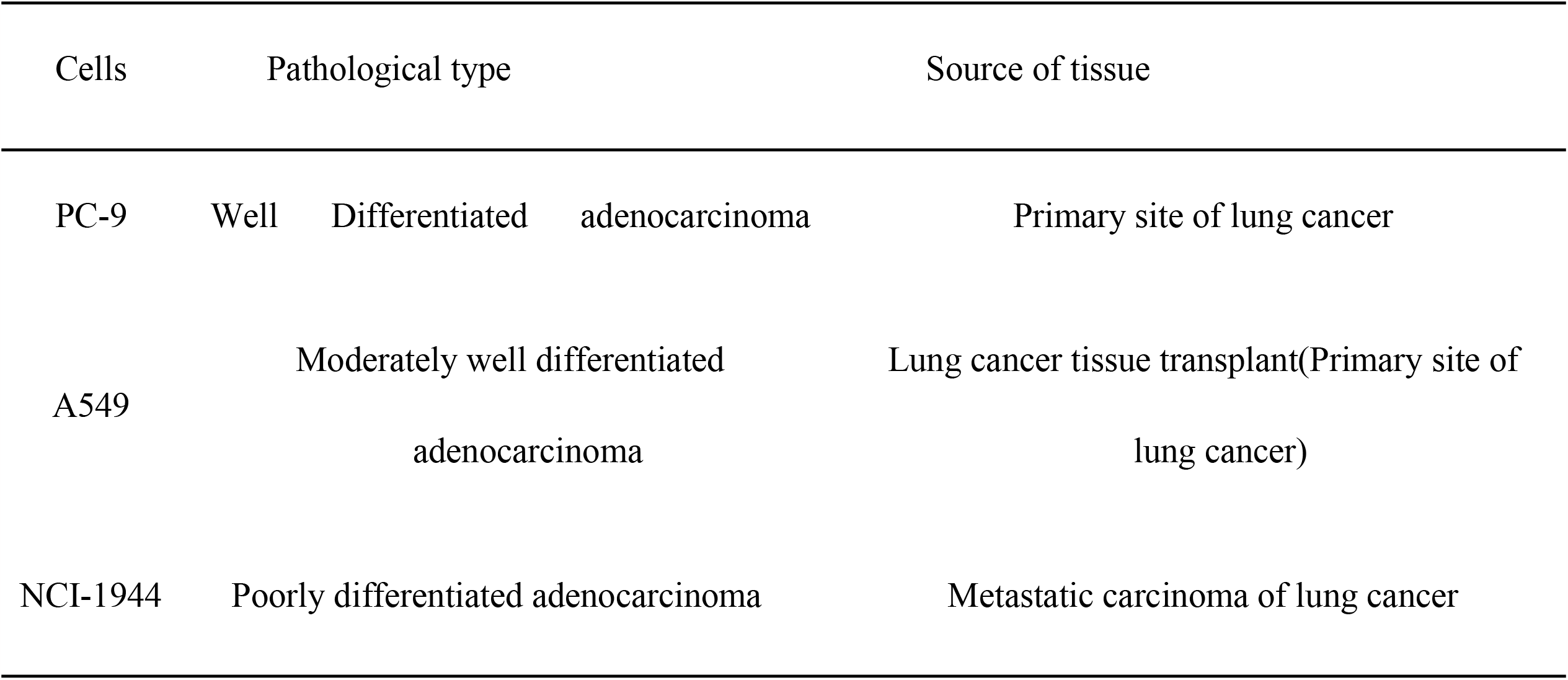
Pathological origin of A549, PC-9 and NCI-H1944 cells.

### Association between overexpression of ALOX15 and proliferation and migration of LUAD

#### Overexpression of ALOX15 inhibits the proliferation of LUAD cells

Through the above experiments, we confirmed that the downregulation of ALOX15 expression was different in LUAD with different degrees of differentiation, and then we examined whether Overexpression of ALOX15 influenced the proliferation of lung adenocarcinoma. We stably transfected lentivirus overexpression of ALOX15 into LUAD cell lines PC-9 and A549 to form stable lines and observed increased expression of ALOX15 compared with normal cells (Fig5. AB). The results of CCK-8 kit showed that the overexpression of ALOX15 in two different LUAD cell lines inhibited the proliferation of cancer cells, and the upregulation of ALOX15 on LUAD cells was the highest at 36h (Fig5.CD).

**Fig 5.**
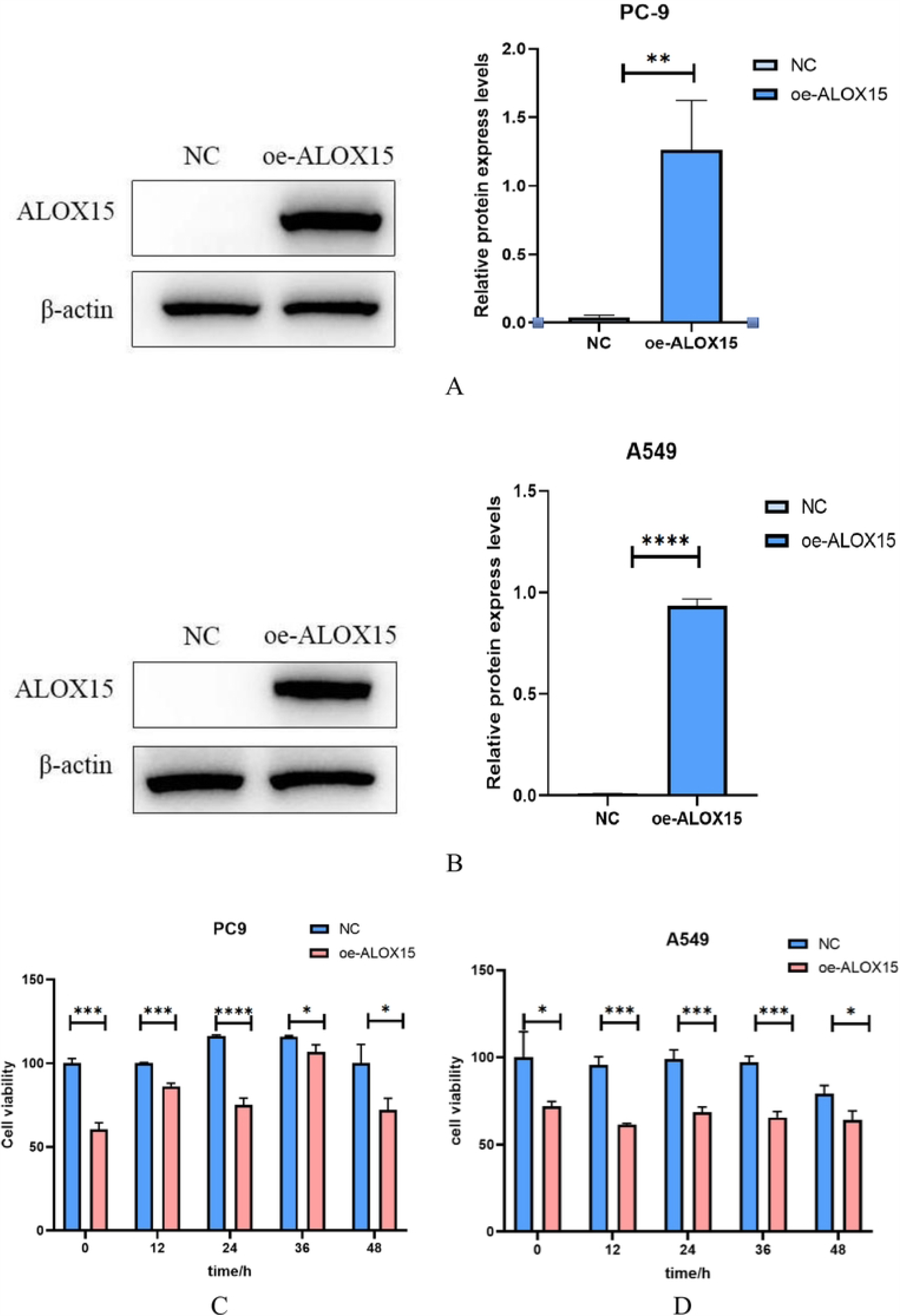
ALOX15 expression levels in different LUAD stable strains overexpressing ALOX15. (A)(B). Cell proliferation was measured in normal LUAD cells and ALOX15 overexpressed LUAD cells, ANOVA was analyzed using a t test.(C)(D). *, P<0.05, \*\**p* < 0.01, \*\*\**p* < 0.001, and \*\*\*\**p* < 0.0001.

#### Overexpression of ALOX15 inhibits LUAD cell migration

The potential role of ALOX15 in regulating LUAD cell migration was investigated by scratch assay in normal LUAD cells and overexpressed LUAD cells. The experimental results showed that PC-9 and A549(Fig6. ABCD) cells overexpressed with ALOX15 closed the scratched area more slowly than normal cells.

**Fig 6.**
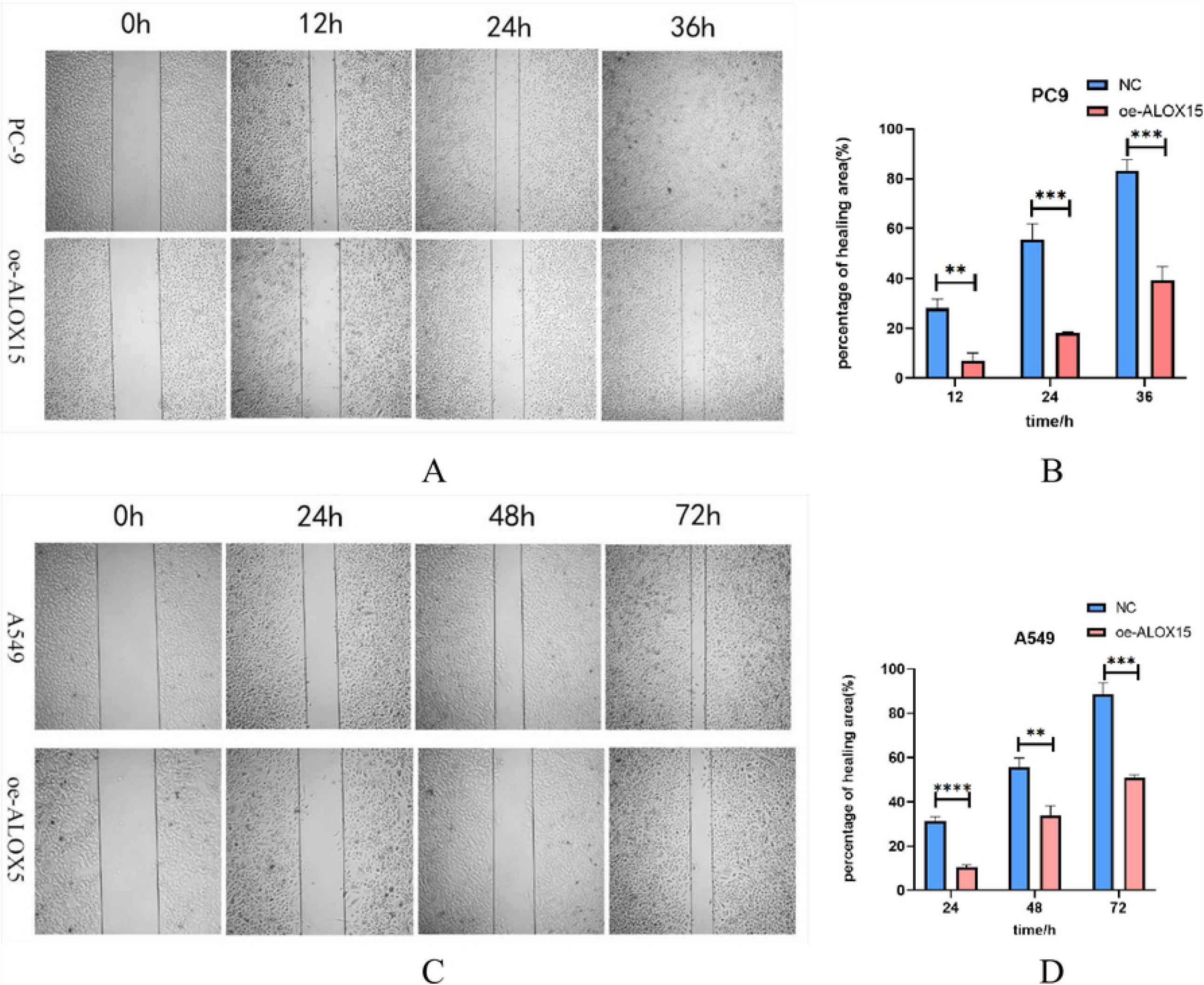
ALOX15 overexpression could inhibit the wound healing ability of LUAD. (A)(B)(C)(D). Wound healing experiments showed that ALOX15 overexpression could inhibit the wound healing ability of LUAD \*\**p* < 0.01, \*\*\**p* < 0.001, and \*\*\*\**p* < 0.0001.

## Discussion

ALOX15 is an inducible and highly regulated enzyme in normal human cells that plays a key role in lipid signaling mediators such as 13(S)-HODE produced by Linoleic acid and Arachidonic acid [21, 22]. Recent studies have found that ALOX15 is downregulated in a variety of major human cancers, including colon, breast, lung, and pancreatic cancers. In this study, we investigated the expression level of lipoxygenase ALOX15 in LUAD and its effect on LUAD. Lung cancer accounts for the second most new cases and the first deaths of all cancers. Conventional treatment for lung cancer includes surgical resection, cytotoxic drugs, and radiation therapy. Immunotherapy therapies are increased with novel targets on cancer. However, lung cancer is still poorly understood with 5-year survival rate of about 15% due to the lack of early diagnosis marker of lung cancer. Therefore, it is more and more necessary to explore the biomarkers affecting the pathogenesis and diagnosis of lung cancer. However, the exact mechanism of ALOX15 in LUAD needs to be further explored in future studies.

Our study provides new evidence for the key role of arachidonic acid 15-lipoxygenase (ALOX15) in LUAD. ALOX15 has been reported to be downregulated in malignant tumors such as lung, colon, esophageal, pancreatic, and breast cancer. TCGA database analysis showed that the expression level of ALOX15 in clinical LUAD cases tended to decrease. At the same time, the expression level of ALOX15 in stage II was significantly lower than that in stage I in clinicopathological stage I-IV, with statistical significance. According to whether the tumor is metastatic and the degree of metastasis, it is found in TNM staging. Stage T refers to carcinoma in situ, stage N refers to lymph node involvement, and stage M refers to distant metastasis. The severity ranges from T to M, ALOX15 showed a trend decrease, especially in the T phase, the expression of ALOX15 was significantly down regulated, with statistical significance. The results of database statistical analysis showed that ALOX15 expression was significantly down-regulated in situ lesions, but we found that ALOX15 expression was also significantly down-regulated and statistically significant through clinical samples of carcinoma in situ and metastatic cancer.

A large body of evidence highlights the critical role of the ALOX15 pathway in regulating tumor cell growth [23]. To further investigate the effect of ALOX15 on LUAD, we collected clinical samples of LUAD. The results of this study showed that the protein expression of ALOX15 gene in lung cancer tissues was mainly distributed in the nucleus, with a small amount of staining in the cytoplasm. The staining of adjacent normal lung tissue was significantly enhanced compared with that of lung cancer tissue. The protein expression of ALOX15 in lung cancer tissue was significantly lower than that in adjacent normal lung tissue, and the expression of ALOX15 in lung cancer was downregulated. It is suggested that the deletion of ALOX15 protein may play an important role in the occurrence of lung cancer. The expression of ALOX15 protein is related to the tissue type and clinical stage of lung cancer, as well as the degree of tissue differentiation and lymph node metastasis.

The difference between the tissues and cells of a tumor and the normal tissues and cells is called the heterogeneity of the tumor, and the degree of differentiation of the tumor refers to the degree of similarity between the tumor cells and tissues and the normal cells and tissues from which the tumor originated. Many studies have shown that the metastasis of cancer is related to the severity of tumor differentiation[24]. The higher the heterogeneity of cancer, the stronger the metastasis and the lower the differentiation degree. Highly and moderately differentiated cancers grow more slowly and are less likely to spread to other parts of the body. In contrast, poorly differentiated and undifferentiated cancers are more aggressive, grow faster and are more likely to spread to other parts of the body. Therefore, in our study of LUAD cell lines with different degrees of differentiation, we found that the expression level of ALOX15 was higher in highly differentiated adenocarcinoma PC9 than in moderately differentiated A549.These studies indicate that the expression level of ALOX15 protein in poorly differentiated lung cancer is lower, indicating that the lower the expression level of ALOX15 protein is, the higher the malignant degree of lung cancer is. By upregulating ALOX15, the proliferation and migration of LUAD cells were inhibited.

Recent studies have shown that the downregulation of ALOX15 expression in colorectal cancer leads to the development of cancer, revealing the high incidence of ALOX15 expression deficiency in human colorectal cancer (close to 100%)[18, 25]. These results are consistent with our results in LUAD. In this study, we investigated the expression level of ALOX15 in LUAD and its effect on LUAD. However, the exact mechanism of ALOX15 in lung adenocarcinoma needs to be further explored in future studies.

## Conclusions

ALOX15 decreased significantly in the early stage of lung adenocarcinoma (LUAD); ALOX15 was down regulated in LUAD with low differentiation and metastasis; The lower the degree of differentiation, the less the expression of ALOX15. The overexpression of ALOX15 in lung adenocarcinoma cells inhibits the proliferation and migration of lung adenocarcinoma. These results indicate that the expression of ALOX15 is closely related to the differentiation, proliferation and metastasis of LUAD. Intervention of ALOX15 may inhibit the development of lung adenocarcinoma, and ALOX15 has a potential biological therapy target function.

## Acknowledgments

Thanks for the support of clinical samples from Department of Pathology, Second Affiliated Hospital of Shandong First Medical University,shandong,China.

## Data Availability Statement

In this paper the genetic and clinical sample analysis data, can be in a public database TCGA (https://portal.gdc.cancer.gov/) to find relevant information.

## Author Contributions

All authors contributed to the study conception and design. Material preparation, data collection and analysis were performed by, Hui Liu, Shupeng Zhang, hongshu Sui,wenwen Sun, Siyu Xuan,minhua Yao, ping Song, peng Qu and yanping Su. The first draft of the manuscript was written by xiaocui Liu,yangyang Tang and all authors commented on previous versions of the manuscript. All authors read and approved the final manuscript.

